# Phantom touch illusion, an unexpected phenomenological effect of tactile gating in the absence of tactile stimulation

**DOI:** 10.1101/2023.02.07.527260

**Authors:** Artur Pilacinski, Marita Metzler, Christian Klaes

## Abstract

We report the presence of a tingling sensation perceived during self touch without physical stimulation. We used immersive virtual reality scenarios in which subjects touched their body using a virtual object. This touch resulted in a tingling sensation corresponding to the location touched on the virtual body. We called it “phantom touch illusion” (PTI). Interestingly the illusion was also present when subjects touched invisible (inferred) parts of their limb. We reason that this PTI results from tactile gating process during self-touch. The reported PTI when touching invisible body parts indicates that tactile gating is not exclusively based on vision, but rather on multi-sensory, top-down input involving body schema. This finding shows that representations of own body are defined top-down, beyond the available sensory information.

## Introduction

You can’t tickle yourself. If you try sliding a finger along your forearm, the tickle sensation will be much weaker than if there was an insect crawling down your skin. This is because the nervous system cancels out the predicted sensory input caused by your own movements [1, 2]. This mechanism is called tactile gating. Tactile gating has been described on neuronal level as downregulation of neuronal firing in response to tactile receptors activation [1, 2, 3]. It affects several levels of processing of tactile information, including spinal cord, motor, premotor and somatosensory cortex [4]. The involvement of these different levels of processing might reflect the complex nature of tactile gating, linked to multisensory representation of touch based on sensory information, body schema, predictive encoding or active touch [5, 6, 26].

While some authors suggest that tactile gating is sensory-driven [7] recent evidence suggests that it indeed results from predictions encoding self touch. Fuehrer et al. showed that the size of tactile gating depends on whether the touch pattern can be predicted or not [6]. While according to the authors this effect suggests that tactile gating relies on prediction, they also show that it occurs for non-predicted stimuli, suggesting that non-specific attenuation may also be involved [6]. Regardless of specificity, the presence of predictive processing in tactile gating makes one ask what if these predictions are confronted with lack of actual tactile input to attenuate. But what happens with tactile gating if there is no afferent tactile signal? Here we tested this using an immersive virtual reality (VR) scenario in which we asked subjects to self-touch using a virtual stick (Figure 1 A). We were inspired by the phenomenon of “phantom” touch sensation that has been anecdotally reported by people using virtual reality environments [8]. This sensation is only broadly described as a vague feeling of touch when - for example - the user gets in contact with another user’s avatar in a VR environment. We predict that, if this sensation results from predicted touch, it should likewise reflect the phenomenological properties of the neural activity meant to attenuate predicted touch on one’s own skin such as moving together with the visual object.

**Figure 1.**
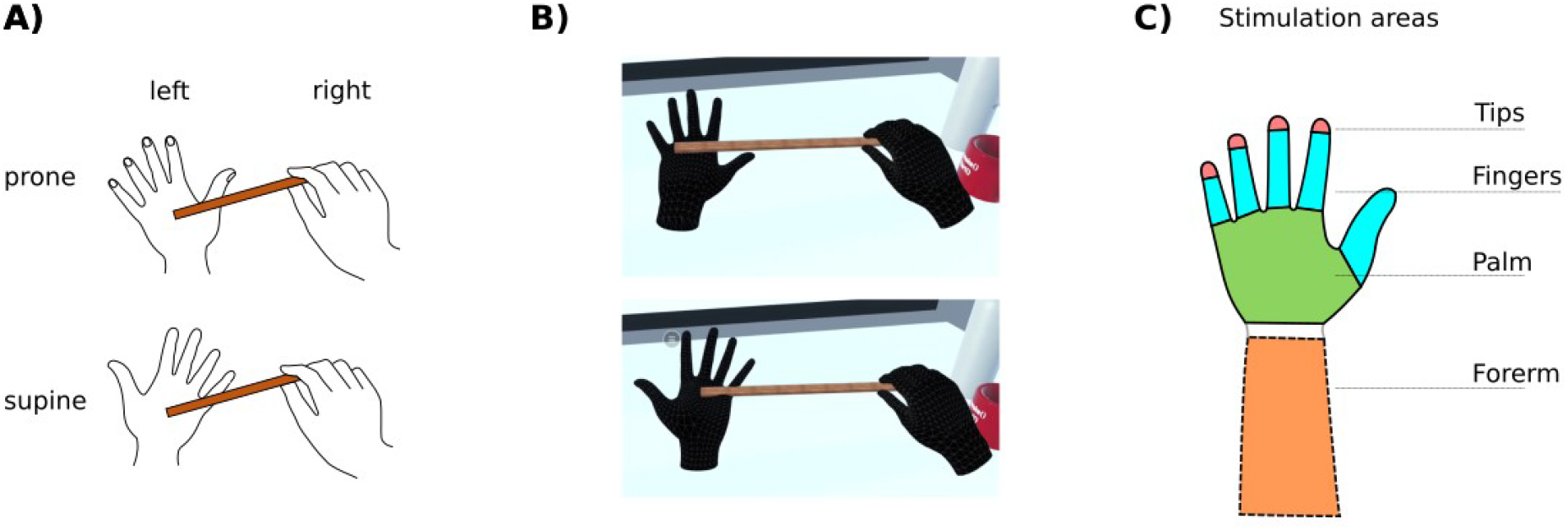
A) Schematic depiction of stimulation with the virtual stick in prone and supine positions. B) A screenshot of the VR stimulation scene in prone and supine positions. C) Stimulation sites: fingertips, fingers (phalanges), palm and forearm. The forearms were invisible to subjects (see panel B).

## Methods

### Phantom Touch Experiment

#### Participants

Thirty-six participants (Fourteen males) took part in the study. The participants were 21-42 years old. The participants had none to little prior experience with virtual reality, no expert knowledge in touch perception, and were completely naive to the experimental hypothesis. The experiment was conducted by three experimenters according to a standard protocol. All procedures were approved by the local ethics committee of the Ruhr-University Bochum and were performed in accordance with relevant ethical guidelines. Subjects provided an informed consent prior to the beginning of the experiment.

#### Experimental setup and virtual reality scenes

We used Unity (Unity Technology, San Francisco, USA) version 2021.3.8f1 and Oculus Quest 2 (Meta, California, USA) head-mounted display (HMD) in hand tracking mode. The scenes took place in a simulated virtual environment where several realistically scaled daily objects were placed on a table in front of the participant (Figure 1 B). Among them there were two sticks, approximately 30 cm long and 2 cm wide. The subjects could only see their virtual hands but not their forearms or other body parts.

#### Procedure

At the beginning, the HMD was put on the subjects and we ensured they could see the scene and their virtual hands clearly. Then, subjects familiarized themselves with the scene such as freely moving and touching the objects on the table. After they appeared acclimatized to the VR environment, the experimental phase began. We asked the subjects to grab one of the virtual sticks by the end with their right hand, and repeatedly stroke their left virtual hand with the stick’s end (See Figure 1B and Supplementary Video 1). We did not further instruct the subjects about any detailed way of stroking the hand such as the exact location to start touching the hand or rhythm. We said that the stick can immerse a bit in the virtual skin, but told them explicitly to be careful to not put the stick through the touched hand and to keep the hands apart (at stick length) and avoid touching their own body at all times. After approximately 30 to 60 seconds we asked the subjects “if they could feel anything”. If they responded negatively, we let them touch their hand with the stick longer and repeated the question after a while. We did not record the exact time of asking these questions nor their number. Importantly, we did not cue the subjects as to what the sensation in the “receiving” hand could be.

If the subjects responded positively, we asked them to describe the feeling and noted down their response. We then asked if the sensation moved consistently with the position of the stick on the skin. Then, we asked the subjects to stroke the middle of the inner side of their palm and rate the intensity of the sensation on a scale from 0 to 10 (0 meant no sensation; we did not further specify what these levels exactly meant). Afterwards we asked them to touch the middle of their fingers and rate the sensation there (on the same scale as above). Lastly, we asked them to rate the sensation’s intensity in the fingertips. Next, we asked subjects to touch the top of their hand and repeat stroking all three parts mentioned earlier. After obtaining ratings, we asked the subject to stroke their (not visible) forearm. We then repeated the procedure for the right hand (with the left hand holding the stick). If subjects spontaneously stroke their hand in a different position than the above order, we did not instruct them to change it, instead we continued the procedure as above until probing all described locations (Figure 1 C).

#### Control experiment

We also tested whether the PTI would happen also when not predicted by tactile gating model, that is without having a visual object touching the hand, for example due to the influence of suggestibility and demand characteristics [9]. For this, we designed a control experiment in which we recreated the basic procedures of the main task albeit without a stimulation to the hand delivered by a realistic object. The procedures mirrored those from the main experiment except from that the subjects were not using a virtual reality headset and, instead of a virtual stick, they held a small laser pointer in their right hand. The laser pointer could project a small (ca. 5mm diameter) point of red light onto the skin of participant’s left hand. There were two conditions: “Laser on”, in which we asked the subjects to stroke their hand with the laser point of light and “Laser off”, in which we asked the subjects to move the pointer as if they were stroking their hand with the point of light. The condition order was counter-balanced and randomized across participants. See Supplement 1 for a detailed description and results.

## Results

All but four of the subjects reported a tactile sensation when touching parts of their hand with a stick. The sensation was usually described as “tingling/static/prickle/electric” or “like wind passing through the hand” (See Figure 2 B and Supplementary Table 1). We call this sensation “phantom touch illusion” (PTI). All subjects that reported this sensation likewise reported that it was consistent with the position of the stick on the avatarized hand. Interestingly, some subjects spontaneously reported they “can feel something” before we asked them for the first time. All subjects described feeling surprised by the sensation caused by PTI. One subject even asked if we were using some device for remote tactile stimulation. The proportion of subjects reporting any sensation (89%) was significantly higher than for control condition without visual stimulation (44%) as evident from proportion test, *Z* = 3.97, *p* < .001 (PTI compared to condition “Laser Off”, see Supplement 1 for details on control experiment). Most importantly, the proportion of subjects consistently reporting “Tingling” responses was significantly higher in the main condition (50%) than in both control conditions: “Laser On” (13%; *Z* = 3.3, *p* < .001) and “Laser Off” (9%; *Z* = 3.62, *p* < .001).

**Figure 2.**
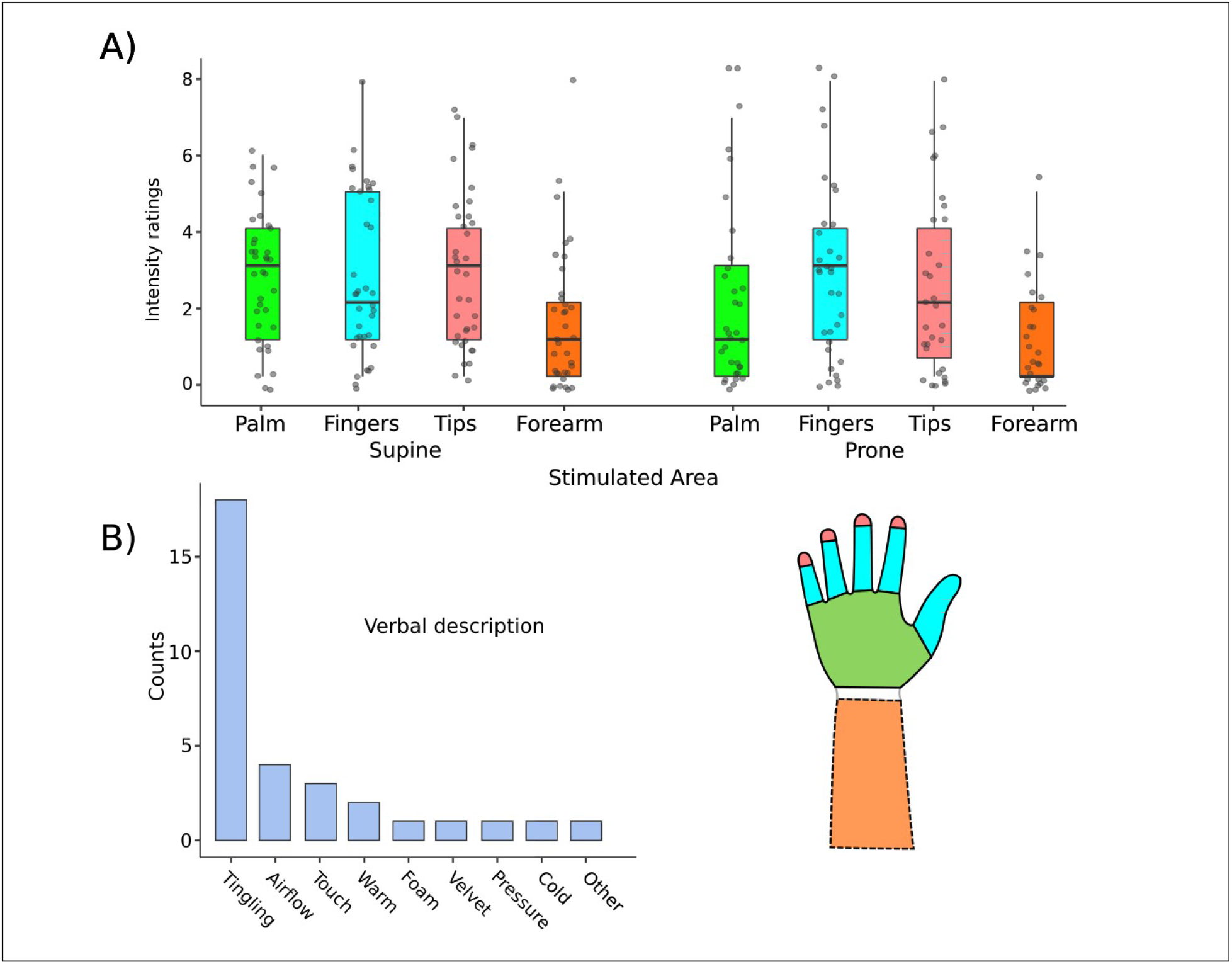
A) Left-hand results of phantom touch intensity ratings across different locations on the hand and the forearm. Box colors represent the relevant parts of the hand as depicted on the right; gray dots represent individual data points; horizontal line represents the median; whiskers represent 1.5 times the interquartile range. Compare Supplementary Figure 1 for the whole range of results across all positions in both hands. B) Frequencies of verbal descriptions of the PTI sensation.

Most importantly, most subjects (Figure 2 A, Supplementary Figure 1; Supplementary Table 1) also reported the sensation to be present when touching their non-visible forearm. Two subjects were visibly startled by this feeling.

The reported intensity of the stimulation differed across subjects (see Figure 2) but in our analysis did not significantly differ across stimulation sites and positions, as shown by repeated-measures non-parametric (Friedman) ANOVA for the left hand (χ^2^[5]=7.83, p=0.16). As we did not control exact stimulation spots, we did not run more detailed tests regarding distinct intensities of the phantom sensation at different spots of the hand surface within each area of interest.

Lastly, several subjects reported PTI when actively touching objects available on the scene (see Supplementary Table 1). Unfortunately, we added this condition only in the course of the experiment and did not consistently measure the strength of the sensation and whether it corresponded to the touching hand parts. We also did not provide a consistent instruction on whether subjects had to just touch or grasp the object which might have further influenced the results (i.e., the number of subjects who reported PTI).

## Discussion

In this study we used immersive virtual reality to simulate scenes of self-touch with a tool without accompanying tactile stimulation. Our subjects reported a tingling sensation at the spots where the virtual stick touched their virtual hand. This happened despite the lack of any physical stimulation. Interestingly, the majority of the subjects reported feeling this phantom sensation when touching their invisible forearm too. We called this sensation “phantom touch illusion” to relate to an apparently similar phenomenon reported in the VR community for social touch [8].

We ruled out that the sensation was caused by heat, airflow or other physical sensation caused by the moving hand by: 1) ensuring separation of both hands by about 30 cm, that is twice the distance at which temperature sensation is typically perceived on the skin in normal conditions [10]; 2) testing the procedure during pilot sessions with a plastic sheet separating both hands which did not affect the sensation.

While our results suggest that touch gating may be predominantly guided by vision because it was induced by a virtual stick and required subjects to look at it, the presence of PTI when subjects touched their invisible forearm is puzzling. It may indicate that tactile gating uses input from other senses than vision, for example proprioception, or top-down input from the body schema. The influence of other modalities and input sources (such as body schema) on tactile gating needs to be investigated.

The very overlapping descriptions of the PTI quality across some of our subjects for both passive (stick) and active (object) touch suggest that PTI does not represent predictions about specific features of object texture or that it does not differ between passive (self) and active (object) touch. Although recent work from Kilteni and Ehrsson [11] suggests that tactile attenuation and gating may be distinct processes aimed at modulating intensity and precision of touch, respectively. It is, however, worth mentioning that although we did not investigate touch quality or precision in detail, we believe there may be subtle differences in tactile gating depending on predicted touch qualities such as suggested by Fuehrer et al. [6]. On the other hand, the same study showed that tactile gating is not limited to predictable stimuli. This lack of prediction-specific gating speaks in favor of the fact that our reported PTI findings may likewise represent stimulus-unspecific cortical events.

An important question remains about the source of neural activity causing PTI. Electrophysiological data shows multi-level processing of tactile gating: cancellation signals are present in the spinal cord and cortical activity in somatosensory, motor and premotor areas [1, 4]. This multi-level processing likely reflects top-down, predictive control of tactile gating processes before movement is initiated [4, 12]. The reported conscious experience of PTI suggests that visual cues for self-generated touch modulate activity in the somatosensory cortex which is the site for tactile sensations [13]. However, the fact that some subjects felt PTI when stimulating their invisible forearm, suggests that its nature may be originating more from top-down modulation of somatosensory cortex through the body schema rather than being purely sensory-driven. Posterior parietal cortex has been often suggested to be a critical site for both the body schema and action predictions [14, 15] and a state comparator across different sensori-motor modalities [16]. As such, it may provide top-down input to the somatosensory cortex in the absence of direct vision of the stimulated skin surfaces.

### Difference to embodiment illusions

While our finding involves feeling touch that is not physical, it does not seem to relate to widely-described embodiment illusions using virtual reality, such as rubber hand illusion [17]. These illusions use simultaneous visual stimulation delivered to an artificial body part (which for example looks like a hand) and tactile stimulation to the user’s own corresponding body part which results in an induced feeling that stimulating the artificial hand is like that of the real one. Interestingly, and in line with our results, such induced embodiment can also be applied to invisible body parts [18]. The main difference of phantom touch to these other embodiment illusions is that PTI did not involve any sort of tactile stimulation that would then be “transferred” to the virtual hand. We can not completely rule out, however, that the phenomenological effects reported in these illusions involve some sort of predictive tactile gating like we believe our PTI results from. This needs more research.

### Open questions

Four subjects did not have PTI, suggesting at least some context specificity that may be due to attentional or other cognitive demands. Our control experiment, together with spontaneous anecdotical reports of PTI in virtual environments do not indicate that the tingling sensation was caused by the experimental procedure. It remains open, however, how top-down cognition influences the specific quality of illusory sensations. Such cognitive influences may be similar to effects reported before, where context seemingly affected object grasping, reflecting qualitative features (temperature) of objects that users grasp in VR [19, 24]. Interestingly, one subject still reported feeling PTI when touching a cup, which indicates there may be differences between active and passive touch as suggested by previous research [11].

Another interesting issue is whether self-reported values reliably represent the perceived intensity of phantom touch and putative differences across areas. While we report lack of differences in reported intensity values across areas, we believe that our experiment was not optimized for answering this question as it lacked an objective control of touch. We believe, nonetheless, that this is worth pursuing further with a more sensitive procedure.

It was recently suggested [9, 20, 21] that embodiment illusions such as the rubber hand may be prone to the influence of demand characteristics of the experimental conditions and phenomenological traits such as suggestibility or phenomenological control. In the case of our experiment, we minimized this confound as we did not provide subjects with any cues about what (especially what touch quality) they should expect and all subjects who had PTI expressed surprise when they felt it. Likewise, the high level of cross-subject congruence in verbal descriptions of the sensation further suggests PTI does not arise from demand characteristics. Such congruence was missing from our “Laser Off” control experiment. It is noteworthy that the actual influence of suggestibility on embodiment illusions has been also challenged [20, 22, 23].

In any case, possible modulatory effects of cognitive demands on PTI, such as mentioned above for temperature perception or sharpness, seem worth investigating.

## Conclusions

We demonstrated an illusory phantom touch sensation emerging during self touch stimulation in the absence of actual tactile stimulus. This PTI was present also when subjects stimulated an invisible, inferred body part. PTI reflects cortical processing of tactile gating resulting in conscious tactile sensation. Moreover, its reported presence in the absence of the vision of the body parts suggests that tactile gating is not just sensory-driven but additionally influenced by the body schema.

## Acknowledgments

This research was supported by the Deutsche Forschungsgemeinschaft (DFG, German Research Foundation) – Project ID 122679504 – SFB 874 and Bial Foundation (Grant: 260/22). The authors would like to thank Gabriella Andrietta, Sophia Kim Bertoni and Tolga Özdemir for help with data collection, and Jon Walbrin, Gavin Buckingham and Pete Lush for discussions.

## Data availability statement

All data generated or analysed during this study are included in this published article [and its supplementary information files]

## Supplementary materials

**Supplementary figure 1.**
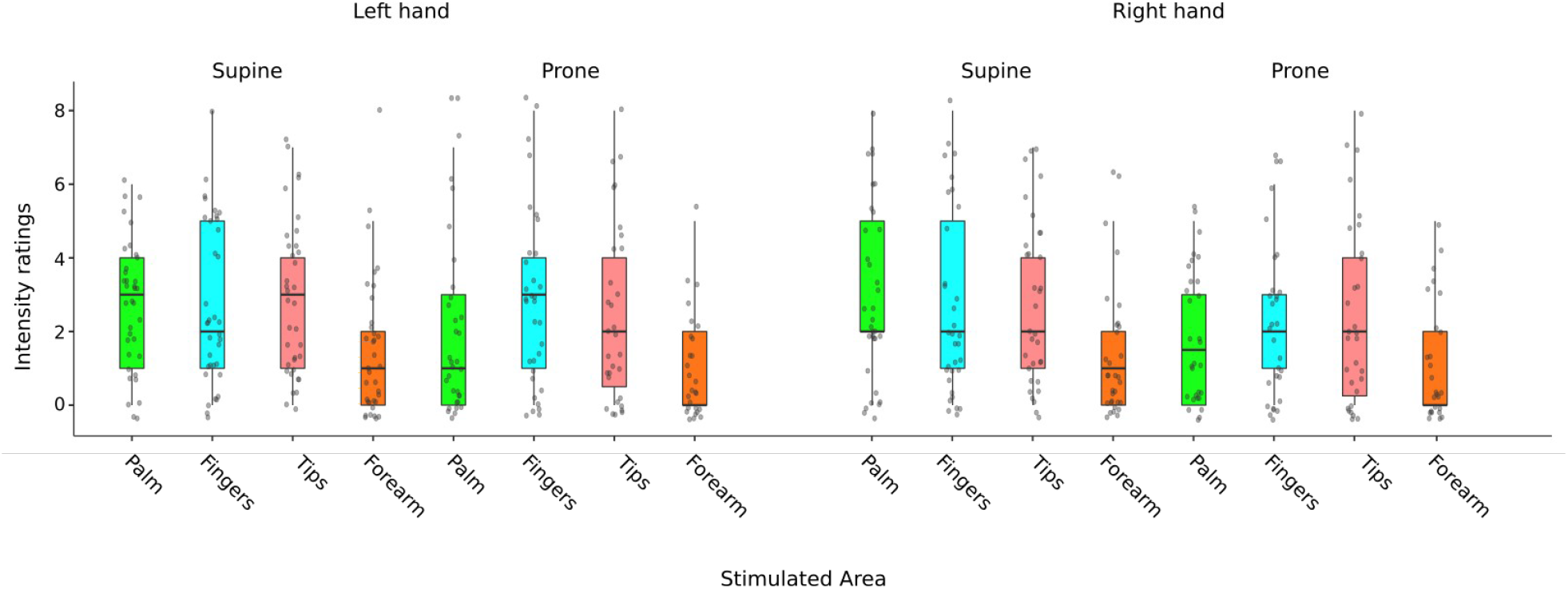
Left- and right-hand results of phantom touch intensity ratings across different locations on the hand and the forearm. Box colors represent the relevant parts of the hand as depicted on the right; horizontal line is the median; gray dots represent individual data points; whiskers represent 1.5 times the interquartile range.

**Supplementary Table 1.**
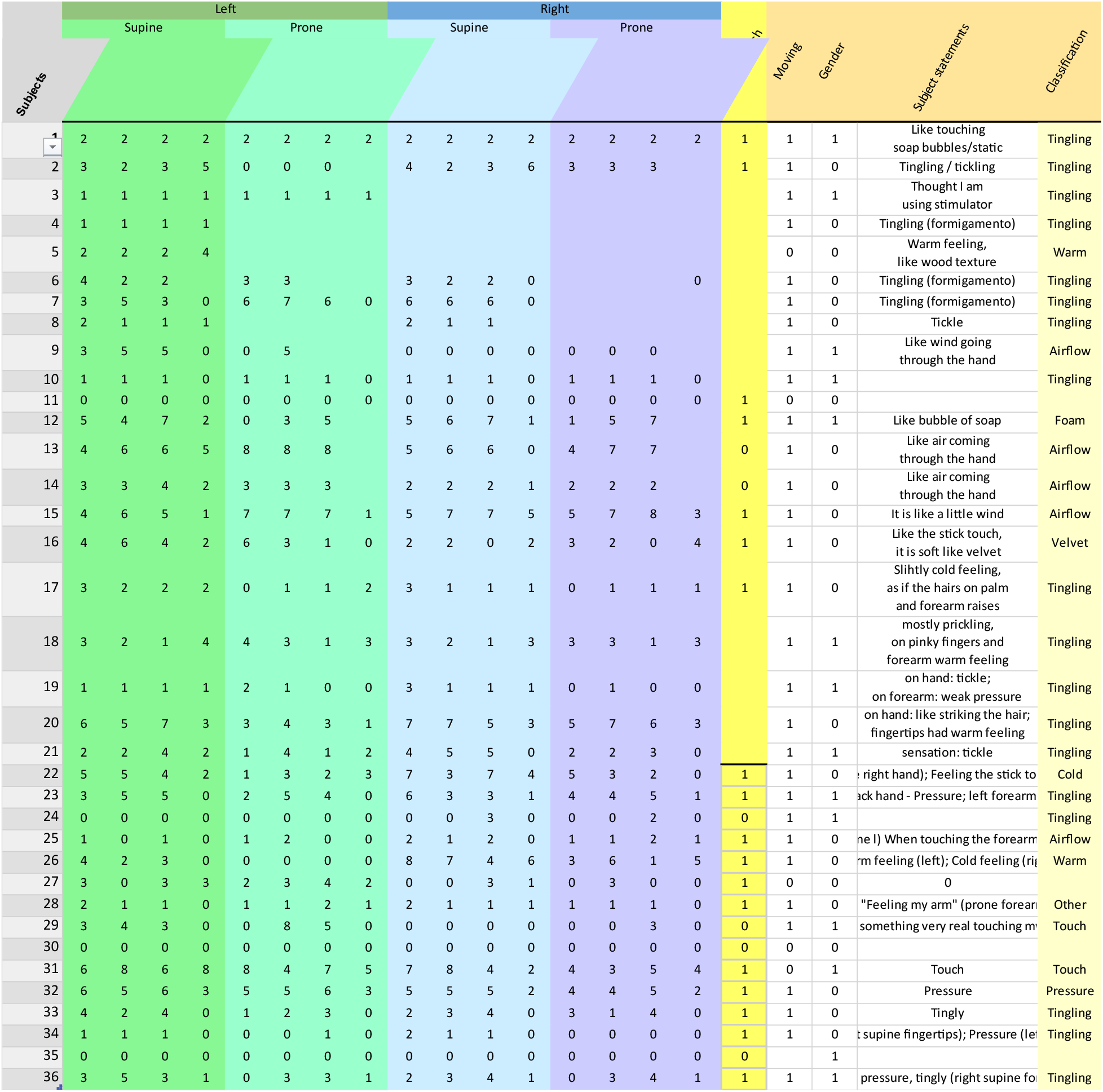
Individual subjects’ ratings of PTI intensity, subject data, specific verbal descriptions and their classifications.

## Supplement 1

### Control experiment

#### Methods

##### Subjects

Thirty-two volunteers (fifteen males), all right-handed participated in the experiment. Fourteen of these subjects also participated in the main VR experiment (the order of main vs. control experiments was randomized and counterbalanced across subjects). All of the subjects were naiive to the experimental hypotheses and had none or little prior experience with VR. All procedures were approved by the local ethics committee of the Ruhr-University Bochum and were performed in accordance with the declaration of Helsinki. Subjects provided an informed consent prior to the beginning of the experiment

##### Procedures

The procedures mirrored those from the main experiment except from that the subjects were not using a virtual reality headset and, instead of a virtual stick, they held a small laser pointer in their right hand. The laser pointer could project a small (ca. 5mm diameter) point of red light onto the skin of participant’s left hand. There were two conditions: “Laser on”, in which we asked the subjects to stroke their hand with the laser point of light and “Laser off”, in which we asked the subjects to move the pointer as if they were stroking their hand with the point of light. These conditions aimed to test subjects’ tendency to 1) report “tingling” in the case a visual stimulation was delivered to the hand previously demonstrated to produce thermal sensation [1] or 2) report tactile stimulation in the case no visual stimulation was delivered to the hand but hand was merely attended, in a way similar to reported by Cataldo et al. [2]. The condition order was counter-balanced and randomized across participants.

### Results

In the “Laser on” condition, 29 out of 32 subjects reported having either a thermal or a tactile illusion, in accordance with previous studies. Of these, 13 reported feeling “warmth” and 4 “tingling” where the laser spot illuminated their skin (See Supplementary Figure 2). In contrast, in the “Laser off” condition, 14 out of 32 subjects reported feeling any sensation, without a clearly dominant quality with just 3 subjects reporting “tingling” (Supplementary Figure 2). Results of the proportion test indicated that there is a significant difference between “Laser off” and “Laser on” in number of reported perceptions *Z* = 3.99, *p* < .001.

---

**Supplementary Figure 2.**
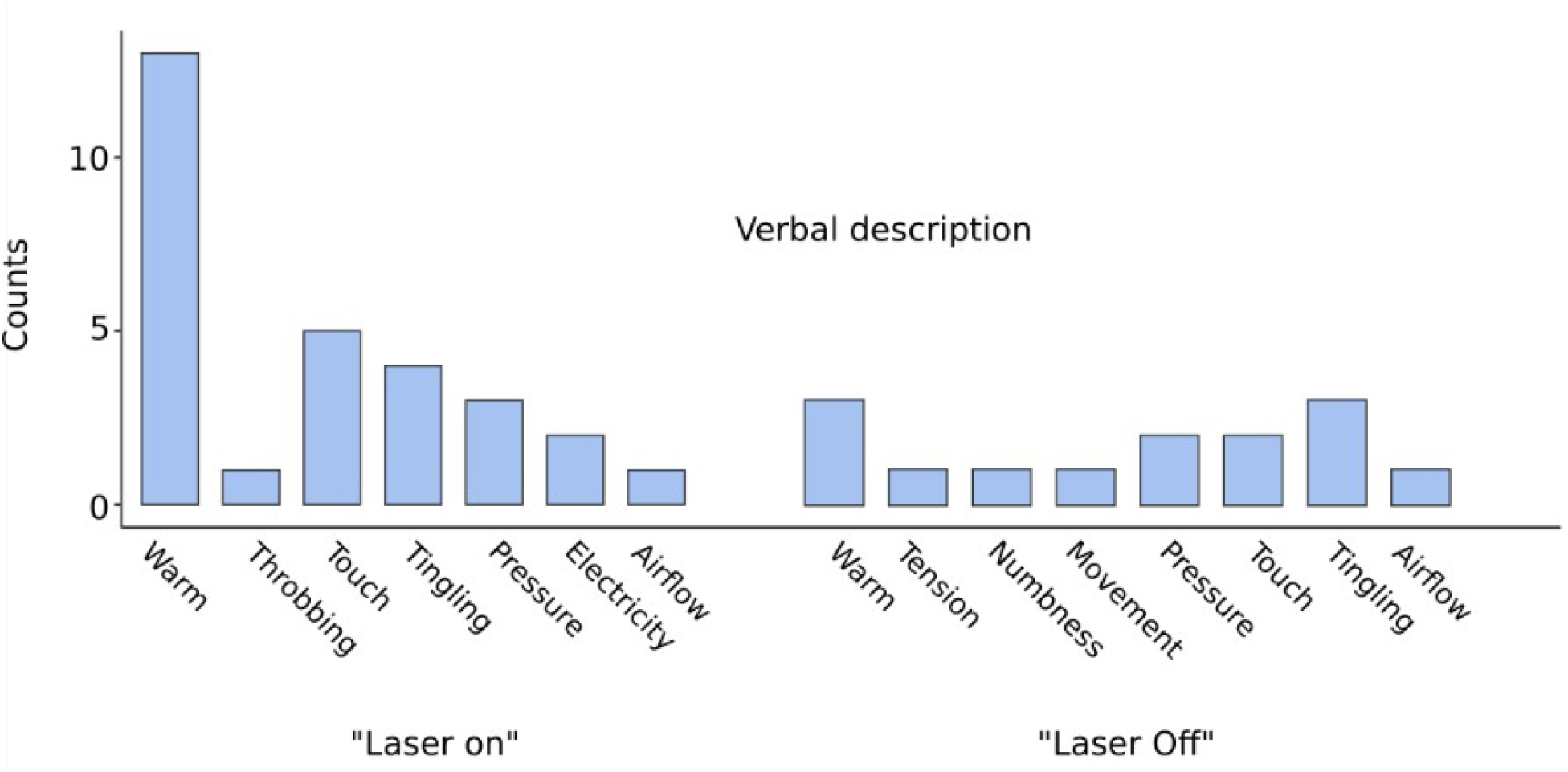
Frequencies of verbal descriptions of the skin sensations in the control experiment for “Laser On” and “Laser Off” conditions.

